# A Spiking Neural Network with Continuous Local Learning for Robust Online Brain Machine Interface

**DOI:** 10.1101/2023.08.16.553602

**Authors:** Elijah A. Taeckens, Sahil Shah

## Abstract

**Objective:** Spiking neural networks (SNNs) are powerful tools that are well suited for brain machine interfaces (BMI) due to their similarity to biological neural systems and computational efficiency. They have shown comparable accuracy to state-of-the-art methods, but current training methods require large amounts of memory, and they cannot be trained on a continuous input stream without pausing periodically to perform backpropagation. An ideal BMI should be capable training continuously without interruption to minimize disruption to the user and adapt to changing neural environments.

**Approach:** We propose a continuous SNN weight update algorithm that can be trained to perform regression learning with no need for storing past spiking events in memory. As a result, the amount of memory needed for training is constant regardless of the input duration. We evaluate the accuracy of the network on recordings of neural data taken from the premotor cortex of a primate performing reaching tasks. Additionally, we evaluate the SNN in a simulated closed loop environment and observe its ability to adapt to sudden changes in the input neural structure.

**Main results:** The continuous learning SNN achieves the same peak correlation (*ρ* = 0.7) as existing SNN training methods when trained offline on real neural data while reducing the total memory usage by 92%. Additionally, it matches state-of-the-art accuracy in a closed loop environment, demonstrates adaptability when subjected to multiple types of neural input disruptions, and is capable of being trained online without any prior offline training.

**Significance:** This work presents a neural decoding algorithm that can be trained rapidly in a closed loop setting. The algorithm increases the speed of acclimating a new user to the system and also can adapt to sudden changes in neural behavior with minimal disruption to the user.

## 1. Introduction

Brain machine interfaces (BMI) are systems that process neural signals from the human brain and use the processed signals to control external devices, such as prosthetic limbs [1, 2]. These technologies can be very beneficial to users who have lost limbs or suffered spinal cord injuries by allowing them to have improved mobility and reduced dependence on caretakers. Surveys have indicated that paralyzed patients have generally positive attitudes towards BMIs [3, 4].

BMI systems consists of three major parts: signal acquisition and processing, feature decoding, and device output [5–7], as shown in Fig. 1. The best results in kinematic decoding are typically achieved when signal acquisition is performed by an implanted electrode the brain, which allows enough spatial resolution to distinguish the activity of individual neurons [8]. After signal acquisition, there exist a variety of different decoding algorithms that can be used to extract kinematic information from the neural signals. The Kalman filter is often considered the gold standard in neural decoding, having demonstrated superior accuracy to linear decoders in several studies [9, 10], but machine learning techniques have also recently shown promise, especially recurrent networks such as LSTMs [11, 12]. However, these recurrent networks often take several epochs to train, which can require large amounts of memory [13].

**Figure 1.**
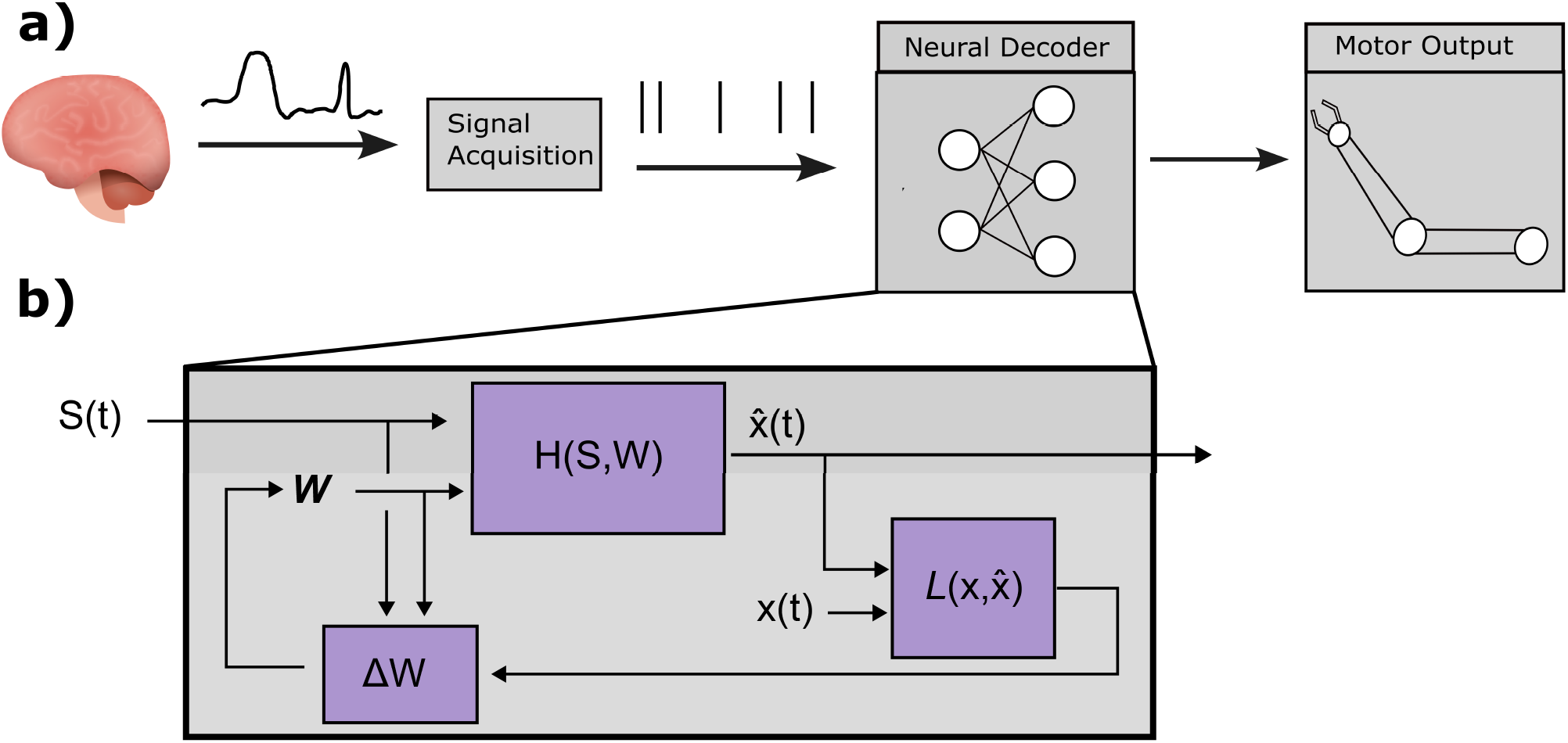
a)An overview of the main components present in a BMI system. A BMI begins with signal acquisition, in which raw voltage readings from electrodes in the brain are recorded and organized into spike sequences via threshold encoding and spike sorting algorithms. These spike sequences are then sent to a decoder, which extracts kinematic information from the neural data. Predicted kinematics are then sent to a motor output, such as a prosthetic limb. b) A more detailed view of the systems present in a neural decoder with continuous learning. The decoder consists of a function which takes neural spiking data *S*(*t*) and a matrix of weights *W* as inputs. The decoder function outputs a predicted kinematic state, 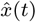. In BMI operation, 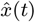 is used to control a motor output. During training, the predicted kinematics can be compared with ground truth kinematic data *x*(*t*) using a loss function, 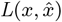. The loss is used to update the weight values.

Spiking neural networks (SNNs) are a type of artificial network that are particularly natural to apply in a BMI application due to their biological nature. SNNs propagate information in the form of discrete sequences of events called spike trains, and process these spike trains using models that closely mimic the action potentials that drive information flow in real biological neurons [14, 15]. This makes them well suited for BMI tasks since the input data from the electrodes is already in the form of biological spikes, so no additional pre-processing is needed. Additionally, SNNs have been shown to be very power efficient due to the sparse nature of the spiking data [16–18], which can be critical for implanted chips, where power dissipation must not exceed 40 *mW/cm*^2^ [19].

Previous work has explored the use of SNNs for neural decoding [20–23]. In [22], authors demonstrated an SNN that could replicate the behavior of a Kalman filter using the Neural Engineering Framework. More recent work has shown that SNNs trained with surrogate gradient descent can also provide highly accurate results. In [20] authors demonstrated that SNNs can achieve superior performance to Kalman filters and artificial neural networks for decoding finger velocity, while [23] showed an SNN decoder with comparable accuracy to an LSTM for decoding arm velocity or body position. However, there are still several challenges that need to be addressed before SNNs can become viable decoding options for a robust and accurate BMI. In previous studies, the SNNs were trained in parallel using short training samples recorded at different points in time. This technique, known as batching, is commonly used to make machine learning algorithms more robust [24], but it is less practical for real world BMI use, in which the system would ideally be trained on chronologically sequential data as the user learns to use it. These SNNs also require more memory as the duration of training data increases, since they must store past spiking events to perform backpropagation. This large memory storage during the training process is not practical for a hardware implementation, which is one of the main motivations for developing SNN neural decoders.

Additionally, SNN decoders have not been tested in an online BMI setting, in which the user attempts to use the BMI in real time and receives information about the output of the BMI. Neural decoders trained offline can have quite different results when applied an online setting due to the adjustments made by the user in response to feedback from the BMI system [25–27]. Even decoders that perform well initially in an online setting may see a decrease in performance over time due to changes in the structure of nearby neurons or sudden loss of neurons [28,29]. Additionally, current online decoding methods often require a period of offline training first, in which the user imagines performing movements [27]. A BMI that can be trained entirely online would require no need for imagined movement on behalf of the user, potentially lowering the difficulty for new users to acclimate to a BMI.

This work presents a spiking neural network for neural decoding that can be trained continuously without the need to pause training and perform backpropagation. Unlike previous continuous learning methods, this novel algorithm is specifically designed for predicting continuous functions such as velocity. There are two main aims of this work: first, to show that the continuous learning SNN decoder can achieve comparable accuracy to existing methods in an offline decoding setting with reduced memory consumption; and second, to show that the continuous learning SNN is capable of quickly adapting to sudden changes in neural behavior and can also learn entirely online with no prior training. To address the first aim, we train the continuous learning algorithm on two different datasets consisting of neural and kinematic data from non-human primates performing reaching tasks [30, 31]. To address the second aim, we train the continous learning SNN in a simulated closed loop environment and evaluate its performance after disruptions to the neural inputs. For both experiments, we compare the performance of the continuous learning SNN with existing SNN decoders, Kalman filter decoders, and a LSTM decoder.

The structure of this paper is outlined as follows: Section 2 presents the architecture of the Spiking Neural Network (SNN), detailing its components and learning algorithms. Within this section, Subsection 2.4 provides an overview of the standard learning rules applied in SNN training, while Subsection 2.5 introduces innovative weight update mechanisms devised for regression tasks in SNNs, facilitating continuous learning. Section 3 details the experimental setup and benchmarks various algorithms. More precisely, Subsection 3.1 discusses the methodologies and findings from experiments with offline datasets, and Subsection 3.2 delineates the closed-loop experimental protocols and their outcomes. Finally, Section 4 reflects on the implications of the proposed continuous learning algorithm and proposes avenues for future research.

## 2. Spiking Neural Network Design

### 2.1. Neuron and Synapse Models

SNNs take as input a sequence of discrete events known as a spike train, defined formally as *S*(*t*) = Σ_*kϵC δ*_(*t − t*^(*k*)^), where C is the set of all events and t^(*k*)^ represents the timing of event k in C. SNN layers consist of neuron models that perform operations on the spiking input and produce their own spike trains that propagate on to the next layer. There are a variety of neuron models, all of which seek to imitate the action potentials found in biological neurons. For this study, we use the leaky integrate-and-fire neuron model [32] due to its relative simplicity to implement, but the work can easily be generalized to other models.

The LIF neuron maintains two internal values: the synaptic current, I(t), and the membrane potential, U(t). The synaptic current and membrane potential of the i-th neuron in a layer L in the network are governed by following:

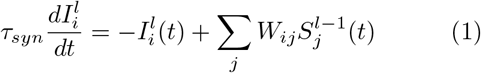

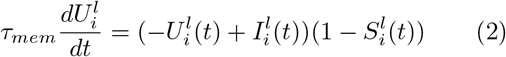

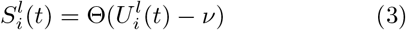

where W represents the matrix of weights applied to the input spikes and *τ*_*syn*_ and *τ*_*mem*_ are time constants [33]. The neuron emits a spike whenever the membrane potential, U(t), crosses a threshold value, at which point the membrane potential is reset to 0.

### 2.2. SNN Architecture

The SNN used in this study consists of an input layer of 46 input neurons, two hidden layers of 65 and 40 neurons, respectively, and a readout layer with two output neurons. The sizes of the hidden layers and other parameters were optimized using Bayesian Optimization [34]. A graphical representation of the SNN architecture is shown in Fig. 2.

**Figure 2.**
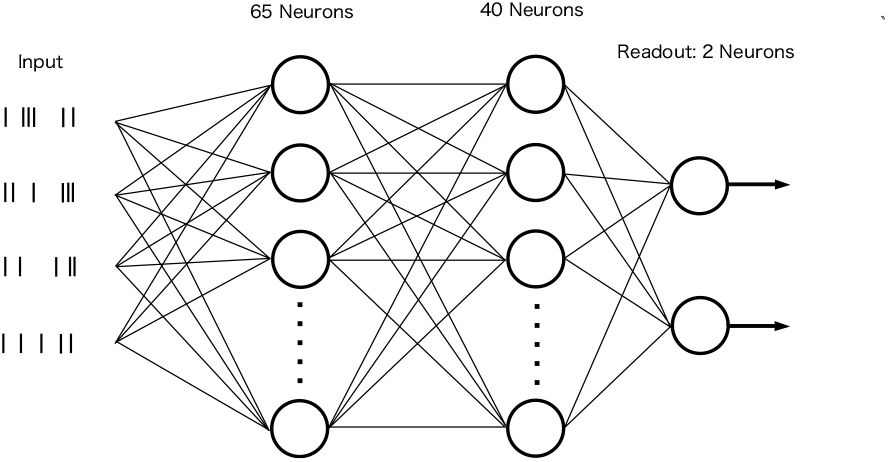
A graphical depiction of the spiking neural network architecture described below. The SNN consists of two hidden layers consisting of 65 and 40 neurons. The output readout layer consists of two neurons corresponding to the velocity in x and y directions.

### 2.3. Readout Function

SNNs are typically used either for classification tasks, in which the readout layer consists of several spiking neurons and the output is determined by selecting the neuron with the highest spiking rate [35, 36], or for modelling spike rates [37]. Neural decoders, however, are designed for regression tasks, in which the decoder outputs continuous real-valued functions. For this study, we focus on the task of predicting 2D velocity of an arm controlling a computer cursor. This task was selected because it is one of the most widely studied BMI tasks [10,38] with several public data sets containing neural and kinematic data [30, 31]. Cursor control has the practical application of enabling BMI users to navigate a computer, granting them greater autonomy. To enable 2D cursors control with a SNN, the SNN output must be modified to produce two continuous real-values functions corresponding to the x and y components of predicted arm velocity. SNN decoders can be adapted to produce these functions by adding a readout layer consisting of two leaky-integrate neurons that never emit spikes [20, 23], where the output is given by the neuron’s membrane potential. This method of computing the output has several desirable properties, such as the fact that the membrane potential is a continuous and differentiable function, so there will be no unnatural instantaneous changes in the predicted velocity. The k-th readout value 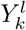 for hidden layer *l* is given by:

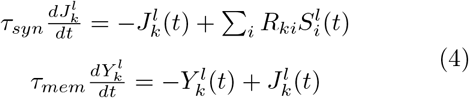

where *R*_*ki*_ represents a fixed readout weight between the i-th hidden layer neuron and k-th readout neuron.

### 2.4. Learning Rules for Training Spiking Neural Network

Traditionally, spiking neural networks have been trained using unsupervised learning rules such as Spike Timing-Dependent Plasticity [39]. However, neural decoders such as Kalman filters and machine learning models are typically trained using supervised learning to extract kinematic information from the neural data [11]. Spiking neural networks can be trained in a supervised manner using surrogate gradient descent, a modified version of gradient descent that approximates gradients for the non-differentiable spiking output of SNNs [33]. However, the calculation of these gradients typically involves backpropagation through both the layers of the network and backpropagation through time to account for the effect of previous spiking events on the output. This requires a large amount of memory that is prohibitive for use in a continuous learning setting. In [40], a technique known as deep continuous local learning (Decolle) is introduced that addresses the issues of backpropagation through both space and time. Local learning removes the need for backpropagation through the neural network by assigning a local readout layer with fixed weights to each hidden layer in the network. The general formula for determining updating weights via local learning is given by

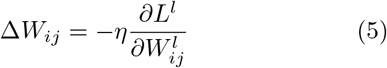

where *η* is the learning rate and *L*^*l*^ is the loss function for the readout of layer *l*. When using mean squared error loss, the loss function is given by:

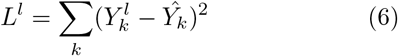

where 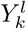 is the kth readout value and 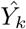 is the corresponding target value. The partial derivative in the weight update equation is then given by:

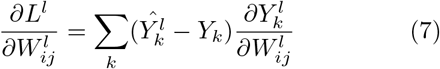

Since 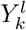 is a function of the output spikes 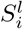 of hidden layer *l*, 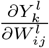 can be expressed in terms of the derivative of 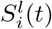 with respect to the weights. Due to the discontinuous nature of spike trains, the derivative of the spiking function, 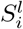, is not well defined. To fix this issue, the heaviside step function 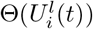 is replaced with a differentiable function 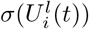 that closely approximates it to perform surrogate gradient descent, as shown in [33]. With this adjustment, the derivative of the spiking output can be expressed as:

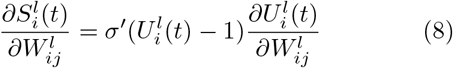

To find 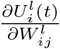 without using backpropagation though time, the equations for I(t) and U(t) are rewritten using the following equivalent expressions:

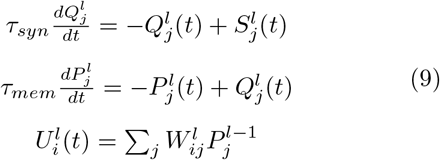

Here, the functions *Q*(*t*) and*P* (*t*) are simply used as a way of rewriting equation 2 in a way that can be differentiated without dependence on past events. *Q*_*j*_(*t*) increases every time neuron *j* emits a spike and decreases exponentially otherwise, while 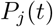 increases at a decreasing rate after each spike. 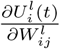 is now simply equal to 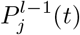.

### 2.5. Novel Learning Rules for Regression

The standard Decolle learning rule works well for classification tasks or simple regressions, but it is not well suited for the task of neural decoding, as we shown in section 3.1.2. This is because the readout function for the Decolle algorithm is a simple linear sum of the spiking output 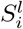. Due to the sparse nature of neural spiking data, this readout function results in highly discontinuous outputs. Predicting velocity for control of a prosthetic requires a system that will preserve important properties of velocity, such as continuity and differentiability, to ensure smooth control. Our readout function, described in section 2.3, uses integration with decay to produce an output that satisfies these properties. This means that the output at any time is a function of previous spiking inputs, not just the current spiking inputs. In order to properly calculate the weight updates, we need to find the derivative of the output with respect to all past spiking outputs. This is distinct from the Decolle algorithm, where only the derivative with respect to the current spiking output is needed, and necessitates a new learning rule. We describe this novel learning rule below.

To find the derivative of the output function 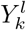 with respect to the weights, it will be useful to rewrite 4 as the convolution of 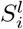 with a kernel function, K(t), as shown in [41]:

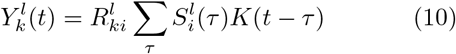

where K(t) is given by

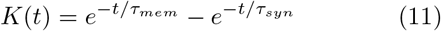

Using the property of derivatives of convolutions, it can be shown that

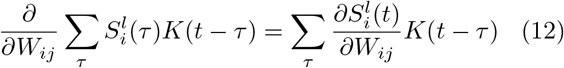

where 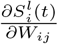 is defined in the previous section by equations 8 and 9.

The expression on the left of equation 12 is mathematically equivalent to the following two differential equations:

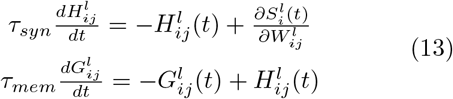

These differential equations integrate 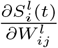, rather than 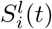. The values *H*_*i*_*j*^*l*^ and *G*_*i*_*j*^*l*^ can be updated at each time step, allowing the algorithm to incorporate the effect of past spikes on the output without actually storing those spikes in memory. The complete derivative of the readout value is then given by:

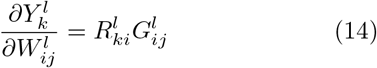

Recall that 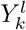 is the output of layer l and is given by 4. Equations 13 and 14 rewrite this equation to allow for calculating the derivative.

Putting all of these together yields the full weight update equation:

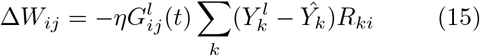

There are several equations used in to calculate the total weight update equation. Table 1 defines the variables and constants used as part of the weight update algorithm and the role of each variable.

**Table 1.** Definition of variables used for learning.

| Variable | Function |
| --- | --- |
| $I_i^l(t)$ | The synapse current of a neuron in layer l. It adds the weighted sum of the spikes from the previous layer with a decay controlled by $\tau_{syn}$. |
| $U_i^l(t)$ | The membrane potential of a neuron. This value integrates the weighted sum of spikes from the previous layer with a decay controlled by $\tau_{mem}$. |
| $W_{ij}^l$ | The weight connecting neuron j in layer l-1 to neuron I in layer l. |
| $S_i^l(t)$ | The spiking output of a neuron in layer l, it emits a spike when $U_i^l(t)$ passes a unit threshold value. |
| $Q_j^l(t),$<br>$P_j^l(t)$ | These integrate the spiking output of an individual neuron in a layer. $U_i^l(t)$ can alternatively be expressed as a weighted some of all $P_j^{l-1}(t)$, where the derivative of $U_i^l(t)$ is given by $P_j^{l-1}(t)$. |
| $Y_k^l(t)$ | The kth readout value for layer l. This value is generated by integrating the weighted sums of spikes output from layer l. |
| $R_{ki}^l$ | The weight connecting the ith neuron in layer l to the kth readout value. |
| $H_{ij}^l(t),$<br>$G_{ij}^l(t)$ | These integrate the derivative of the spiking output $S_i^l(t)$ with respect to weight $W_{ij}^l$, and are used to calculate the derivative of $Y_k^l$. |
| $\tau_{syn},$<br>$\tau_{mem}$ | Constants that control the rate of decay for the synaptic current I(t) and membrane potential U(t), respectively. The decoder performs best when $\tau_{mem} \gg \tau_{syn}$. For offline experiments, $\tau_{mem}$ was in the range of 50-100 ms while $\tau_{syn}$ was between 0.5 and 2 ms. |

This weight update equation accounts for all previous states of both the hidden layer neuron and the readout neurons without requiring any memory overhead to store the historical states of these neurons. It does require supplementary functions in the form of 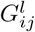 and 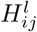, but these only increase the memory complexity to *O*(*NL*), where N is the number of input neurons and L is the input layer size. The memory complexity of BPTT is *O*(*NT* ), where T is the input duration. In BMI applications, L is typically significantly less than T, since SNN performance don’t increase significantly after the layer size increases beyond about 50-60 neurons [23], whereas the number of time steps could easily enter the hundreds of thousands for a continuous experiment that lasts longer than an hour, assuming a neural sampling rate of 100 Hz.

Additionally, because the memory does not have to store past states and produces weight update values at every time step, the learning rule proposed in this work is capable of online continuous learning. When training with backpropagation through time, a SNN must first do a forward pass on a training example, and then stop and backpropagate through the training example to calculate the gradients before moving on to the next example. The continuous learning algorithm proposed in this work evaluates the forward pass and weight updates simultaneously, allowing the weights to be updated at every time step without ever needing to pause the experiment.

## 3. Experimental Procedure and Results

### 3.1. Offline Learning

#### 3.1.1. Evaluation Data

The SNN was trained and tested on two different data sets containing intracortical neural recordings and corresponding kinematic data from non-human primates performing reaching tasks [30, 31, 42, 43]. The first data set, taken from the Collaborative Research in Computational Neuroscience (CRNCS) dataset repository, contained recordings from the premotor cortex of a monkey as it moved a cursor towards an indicated 2 cm by 2 cm target on a 20 cm by 20 cm computer screen. The velocity of the on-screen cursor was sampled at 1000 Hz. Raw neural recordings were performed using a 100 channel Utah electrode array. The neural recordings were preprocessed to form spike trains using standard manual spike sorting techniques [30], yielding 46 independent neurons. Both the neural and kinematic data were placed into 10 ms bins before being passed to the network, with neural data limited to at most 1 spiking event per bin. 5.4% of all spiking events were lost due to the 10 ms binning. The mean inter-spike interval was 175 ms. Each reach lasted up to 1.7 seconds. The second data set, referred to by the authors as “MC Maze”, contains recordings from the motor and premotor cortex of a monkey as it performs reaching tasks while navigating the boundaries of a 2D maze [43]. Neural data was recorded using 2 96-electrode arrays implanted in the PMd and M1 regions. Spike sorting was performed offline to isolate independent neurons; the trial that we used contained 107 neurons in total. Spike times were determined via threshold crossing. Both the neural data and corresponding hand velocity data were placed in 1 ms bins. The authors did not provide information about how many spiking events were lost due to this binning, but the mean inter-spike interval was 221.9 ms. Reaching tasks last up to 600 ms. Decoders were trained on spiking data from 130 ms prior to movement onset to 370 ms after movement onset, per the instructions of the authors of the data set.

#### 3.1.2. Batched Learning Comparison

SNN decoders have traditionally been trained using batched learning methods [20,23]. To ensure a valid comparison with existing decoders, the continuous learning model in this study was also trained using batched learning. For the evaluation of this model, two datasets were used: CR-CNS and MC Maze. The performance of the proposed continuous learning algorithm for SNNs was bench-marked against several established methods: SNN trained using backpropagation-through-time (BPTT), an SNN trained using unmodified DeColle learning algorithm [40], a Kalman filter, and a LSTM decoder. The Kalman filter is often used as a standard benchmark for comparing new BMI decoders [20, 44], while the LSTM was selected due to recent work demonstrating superior accuracy to other methods [11, 45]. Additionally, an SNN trained with unmodified DeColle learning rules described in [40] to demonstrate the necessity of the novel continuous learning algorithm. This Decolle-trained SNN uses a linear readout function and as a result is not expected to perform well for this task; it is included to show the necessity of the continuous learning rule introduced in this paper.

The results for each decoder are shown in Fig. 3 comparing the average peak correlation coefficients achieved by each decoding method across 10 different train-test splits for the CRCNS PMD data set and the MC Maze data set, respectively. All models were trained using 10 different train-test splits for each data set. In each split, 60% of the data was used for training, and 20% for testing. The remaining 20% was isolated from all train test splits and used as validation for optimizing hyperparameters, such as the time constants used to determine the spiking dynamics. The Kalman filter demonstrated higher accuracy when using binned sums of the neural data; a 50 ms sliding window was used to bin the neural data for the CRCNS data set, while a 10 ms sliding window was used for the MC Maze data set. The LSTM also demonstrated higher performance with a 10 ms sliding window for the MC Maze data set.

**Figure 3.**
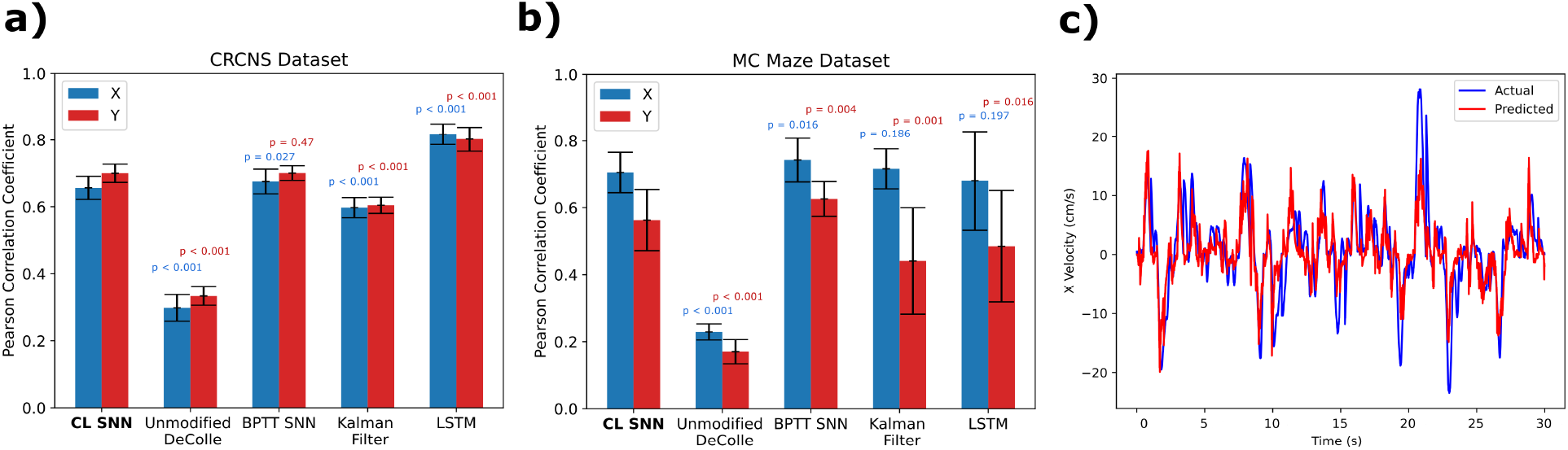
a,b) Comparison of the average correlation coefficients achieved by each decoding method across 10 different train-test splits for the CRCNS PMD data set and the MC Maze data set, respectively. For each train-test split, each model is trained on 60% of the available data, with 20% used for testing. The continuous learning SNN is compared with 4 other decoding methods. The error bars shown for each decoding method are proportional to the sample standard deviation and represent 95% confidence intervals, assuming that the correlations from different splits tend towards a normal distribution. Welch’s t-test was used to determine if the differences in mean correlations were statistically significant. The corresponding p-values are shown above each comparison decoder. The results showed that the continuous learning SNN decoder results were significantly different (*p <* 0.05) in both the x and y direction when compared to the unmodified DeColle, Kalman filter, and LSTM on the CRCNS data set, but only significantly different from the unmodified DeColle on the MC Maze data set. In general, these results indicate that the performance of the continuous SNN on offline datasets is comparable to existing methods. c) A 30 second sample comparing the ground-truth kinematic data with the predicted kinematic data output by continuous learning SNN. The network used to generate this example was trained for 50 epochs.

The barplot in Fig. 3 shows the pearson correlation coefficients which is a square root of coeffecient of determination (*r*^2^). Further the barplot show the error bars that are proportional to standard deviation. Most decoders achieve correlation coefficients between 0.5 and 0.8, with the exception of the unmodified DeColle algorithm, which was not designed specifically for neural decoding. Welch’s t-test was used to determine if the differences in mean correlations were statistically significant; the results showed that the continuous learning SNN decoder results were significantly different in both the x and y direction when compared to the Kalman filter (*p <* 0.05), LSTM (*p <* 0.05), and unmodified Decolle (*p <* 0.001) on the CRCNS data set, but only significantly different from the unmodified DeColle (*p <* 0.01) on the MC Maze data set. The significance value are shown in Fig. 3 at top of the bar plot. These results demonstrate that the continuous learning SNN can achieve comparable performance to an equivalent SNN trained with BPTT and to standard neural decoders.

All decoding methods except the Kalman filter were trained using iterative error-based methods, i.e. each decoder was initialized with random weights and generated predicted velocity outputs using the training neural data as inputs. The mean squared error between the predicted and true velocity was used to update the weights, and the process was repeated for 30 epochs for each decoder. The Kalman filter was trained by fitting measurement and transition matrices to the training data, as described in [10].

#### 3.1.3. Memory Footprint

The continuous learning SNN shows a significantly reduced memory footprint compared to the BPTT learning methods. Reach tasks from the CRCNS data set were typically between 1.0 and 2.0 seconds; for a 50-neuron input with a 1.0 second duration and a batch size of 1, the stochastic gradient descent BPTT algorithm had a minimum memory requirement of 838 KB to store all necessary spiking data and gradients. By comparison, the continuous learning SNN had a minimum memory requirement of just 144 KB to store all necessary information, regardless of the duration of the input. Table 2 compares the amount of memory consumed by each function used during continuous learning and BPTT learning. These values represent lower bounds on the amount of memory used by each system; they are calculated by measuring the size of the arrays instantiated during training multiplied by 4 bytes per floating point value. As a result, the minimum memory requirements are independent of the programming environments used to implement the two learning methods. The actual amount of memory used during implementation cannot easily be compared because BPTT was implemented with the Pytorch machine learning library, which adds a large amount of memory overhead. The majority of the memory used by the BPTT algorithm comes from the need to store the full spiking history of every neuron to calculate gradients. This means that the memory used by BPTT scales linearly with both the input duration and the number of neurons in the network. By contrast, the majority of memory used by the continuous learning SNN is the memory required to store the values *G*_*ij*_ and *H*_*ij*_ for each neural connection. This means that the memory usage of the continuous learning SNN scales linearly with number of input neurons, assuming that the number of hidden neurons stays constant, and is invariant to input duration, as shown in Fig. 4. Even with large input sizes of up to 1000 neurons or electrode input channels, the continuous learning method uses less memory when the training duration is longer than about 500 ms.

**Table 2.**
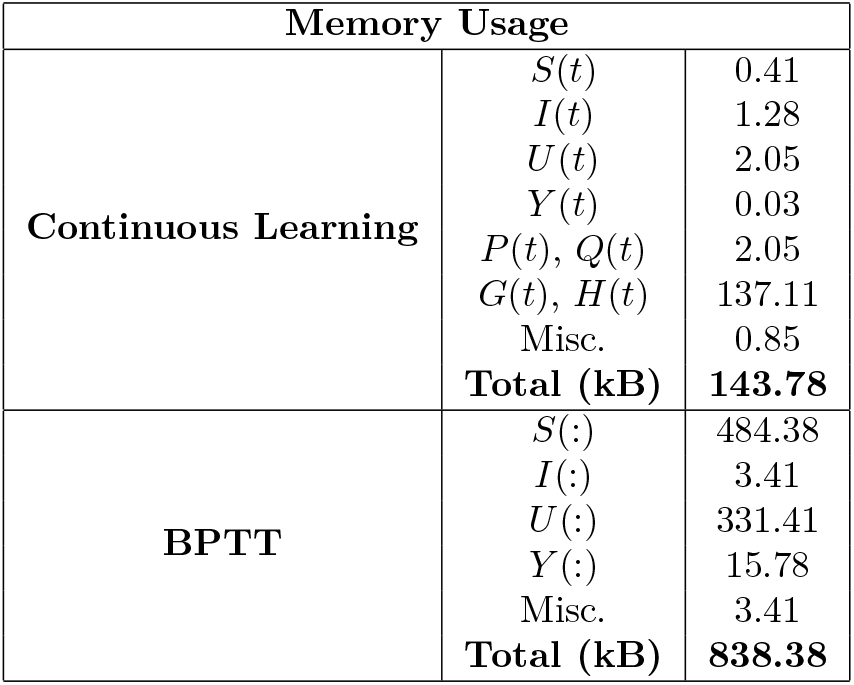
The memory used by each part of the learning process for continuous learning and BPTT during a standard reaching trial (50 input neurons, 1.0 second duration). The continuous learning method uses less overall memory because it only needs to save current values of S(t) and U(t), while BPTT needs to save all past values for these functions.

**Figure 4.**
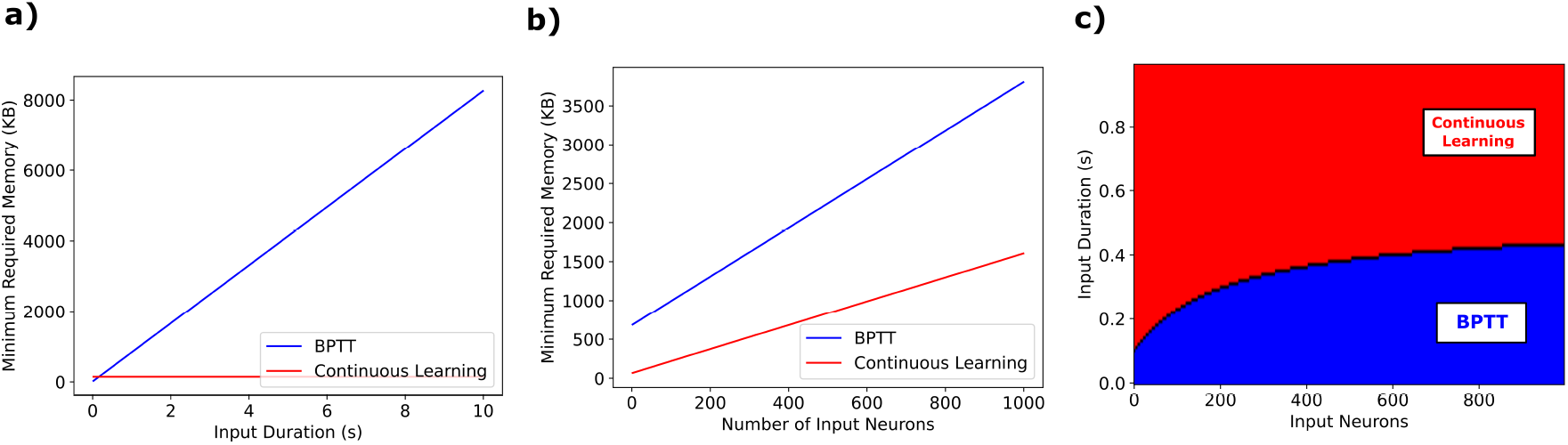
a) Memory complexity for each learning method when the input size is held at a constant 50 neurons. The memory required to perform continuous learning is constant regardless of input duration, whereas the memory required by BPTT scales linearly with input duration. b) The memory required for both learning methods when input duration is held constant at 1.0 seconds, with an input timestep of 10 ms. scales linearly with number of input channels. The slope of the line for continuous learning is determined by the size of the hidden layers, whereas the slope of the continuous learning line is determined by the input duration. c) The conditions under which each learning method requires less memory, assuming 10 ms time step and constant hidden layer size. In the blue region, the BPTT algorithm requires less memory, while in the red region, the continuous learning algorithm requires less memory. The black line represents the boundary where both methods require the same amount of memory. This graph shows that the continuous learning algorithm is more efficient when the input duration is long or when the number of input neurons is small. Even with a large number of input neurons, the continuous learning algorithm is still more efficient if the input duration is any longer than 0.5 seconds. This makes the continuous learning algorithm ideal for training in real time on multiple reaching trials.

#### 3.1.4. Continuous Learning

Current state-of-the-art decoders use batched training, which requires simultaneously training on data that was recorded at different points in time for several epochs. To reduce calibration time and disruptions to the user, a BMI that can learn continuously would make more sense. Surveys conducted on patients with motor impairments show that users prefer BMI systems that offer high accuracy, minimal daily setup, and multifunctionality [3, 46, 47]. To test the continuous learning capabilities of the SNN, we again evaluated it on the premotor cortex dataset, but this time with no batching and without splitting the recording into discrete training examples. Additionally, to test the ability of the network to maintain performance after training stops, we ran the experiment multiple times with different training times. The results show that the SNN converges rapidly, performing most of the learning within the first 300 seconds. The accuracy does not noticeably decline during the testing phase, regardless of when training stops. The peak correlation achieved is less than the peak correlation achieved after multiple epochs of training, but it does achieve the same peak correlation as the batched learning did after a single training epoch. We benchmark the continuous learning performance against that of an SNN trained with BPTT. The BPTT algorithm cannot learn continuously, so we instead allow it to pause and perform backpropagation after each individual reaching task (duration 1.0-1.7 seconds). Both methods eventually achieve the same peak correlation, but the continuous learning mechanism learns more quickly, as shown in Fig. 5. Additionally, BPTT takes longer than indicated in the figure, since it must pause after each reach to perform backpropagation. Backpropagation for one reach takes an average of 35 ms, which introduces a delay in training after hundreds of reaches and also interrupts the user.

**Figure 5.**
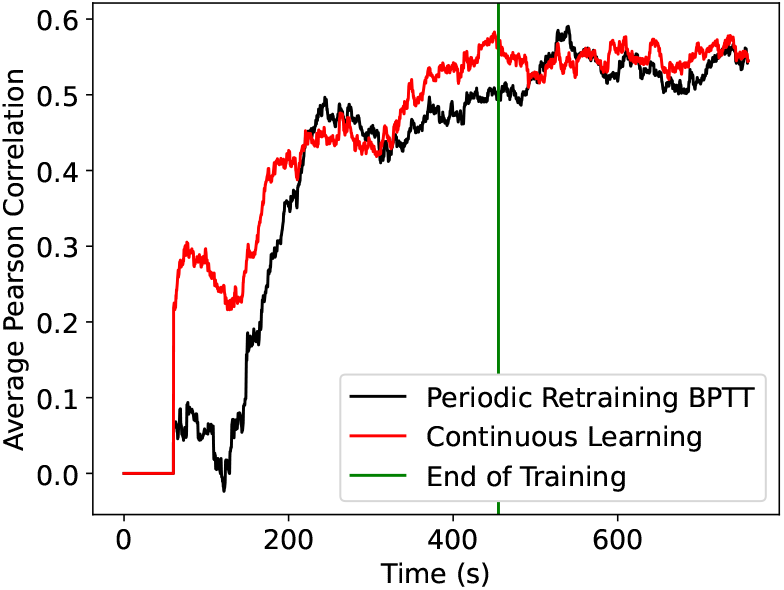
A sliding window average of the Pearson Correlation achieved by two different SNNs trained on a single epoch of reaching data. A sliding window with a 60 second duration is used to smooth out noise in the plots. The X and Y correlations are averaged for each SNN. Although both models eventually reach the same peak correlation, the continuous learning SNN learns more quickly. Both models maintain their performance after training stops. The figure does not include the time it takes for BPTT to pause to perform backpropagation (an average of 35ms for a reach of 1.7 seconds).

### 3.2. Closed Loop Experiments

While offline experiments on pre-recorded data are commonly used to evaluate neural decoder performance, they do not always correlate well with performance in an online, closed-loop setting [25–27]. In a closed loop environment, the output of the BMI is fed directly to the output device, allowing the user to attempt to correct for any errors in the BMI output in real time. This user feedback can have significant impacts on the performance of a BMI system.

#### 3.2.1. Experimental Procedure

Performing a real closed loop experiment on human subjects can be expensive and time consuming. Neural simulations have been shown to produce neural data that more closely mimic the neural responses of real human subjects using a closed loop BMI when compared to real neural data taken from offline experiments [48]. For this work, we adapt the online prosthetic simulator (OPS) described in [26]. The OPS uses the cosine tuning model to determine the spiking rates of neurons [49], in which each neuron has a “preferred direction” and is more likely to fire when the direction of motion is in its preferred direction. The firing rate *λ*_*t*_ of the k-th neuron at time t is given by:

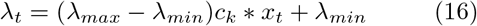

where *λ*_*min*_ and *λ*_*max*_ are the minimum and maximum firing rates, respectively, *c*_*k*_ is a unit vector in the neuron’s preferred direction, and *x*_*t*_ represents the desired acceleration vector at time *t. λ*_*t*_ was constrained to be greater than or equal to zero. Acceleration was used to determine firing rates because the real neural data used in the previous section showed a much stronger correlation with acceleration than with velocity or position. We sampled direction vectors uniformly from the unit circle. Maximum and minimum firing rates were sampled between 40 and 100 and between 0 and 5 spikes per second, respectively. These values were again chosen to match the firing rates found in real neural data, and match closely with values used in previous OPS simulations. In [26], a Poisson distribution with parameter *λ*_*t*_ was used to determine the number of spikes in each time bin. Since SNNs operate on individual spikes, and since Poisson random variables can be represented as an approximation of a sum of many Bernoulli variables, we modeled each neuron as a Bernoulli random variable with spiking probability *p* = *λ*_*t*_ * *t*_*s*_ using a spike sampling period *t*_*s*_ = 10 ms.

The OPS was previously used in [26] to generate simulated neural activity for people performing center-out reaching tasks, with ground truth kinematic data used as the input to generate simulated neural activity. In the absence of real human test subjects, we use a virtual center-out reach task simulation in which targets are placed at random locations a fixed distance away from a central starting point. The ground truth kinematic data that is sent to the OPS is generated by assuming that a real user would always attempt to move in the direction of the target. This assumption has previously been used to train online neural decoders to a high level of accuracy [27]. This, along with constraints on the velocity and acceleration, is used to generate intended kinematic profiles, as shown in Fig. 6a. Target acceleration vectors are then sent to the OPS to generate neural activity, which is in turn decoded by the SNN to produce a predicted velocity and update the position. At the next time step, the new velocity and acceleration vectors go from the current position to the target, as shown in Fig. 6b.

**Figure 6.**
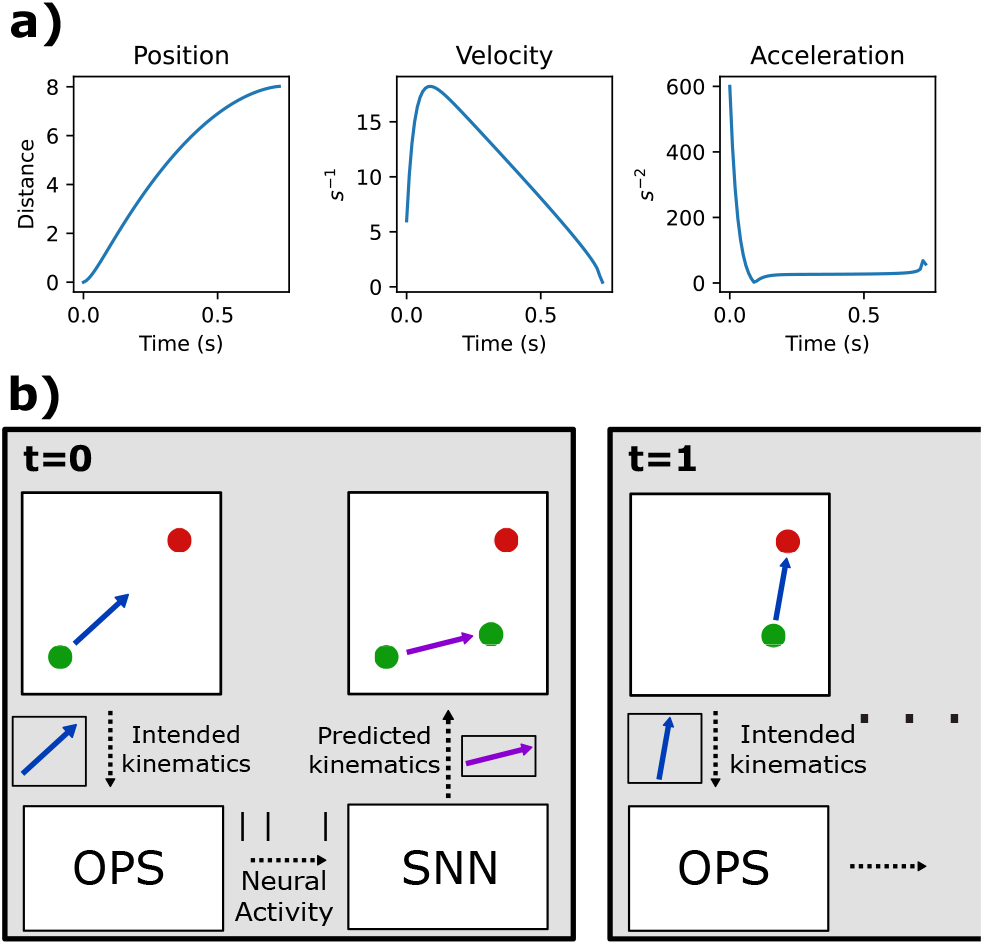
a) A plot of the magnitudes of the intended position, velocity, and acceleration profiles for a sample reach in which the simulator travels along its intended path. The units of time are in seconds (on x-axis), while the distance units are arbitrary. b) Graphical depiction of the feedback system used by the closed loop simulator. The system starts by generating an intended acceleration vector in the direction of the target, which is sent to the OPS to generate simulated neural spikes. The spiking activity is decoded by the SNN to produce a predicted velocity, which is used to update the current position. At the next time step, a new target acceleration is generated based on the updated position.

For all experiments described below, a reach is considered successful if the decoder is able to stay within a unit distance of the target for at least 500 ms (the targets were placed 4 unit distances away from the central starting location). Reach tasks are allowed to proceed for a maximum of 3.0 seconds, after which point they will be considered failed. These values are generally in line with previous center-out reach experiments [27, 42].

#### 3.2.2. Neural Variability and Disruptions

One of the main challenges with using BMIs in a closed loop setting is the decrease in decoding quality that can occur as a result of neural disruptions [28]. We performed several closed loop simulations to evaluate the effects of various neural disruptions on the SNN decoding performance. Prior to the closed loop experiments, the SNN was trained on 400 simulated reaches for 5 epochs using batched learning. The reaches were generated using the online prosthetic simulator described above. To provide standards of comparison for SNN performance, we also trained a ReFIT Kalman Filter, a SNN with BPTT learning, and a LSTM decoder on simulated offline reaches. The ReFIT Kalman filter is a modified Kalman filter decoder designed specifically for closed loop BMI experiments that accounts for the lack of uncertainty in the measured position, since the position is determined by the output of the decoder [27]. The BPTT SNN was included to show how the continuous learning rule affects adaptability. LSTM decoders have shown the ability to surpass the performance of Kalman filters for both offline and online decoding tasks [11, 45], so it provides an additional comparison standard for the current state-of-the-art. The Kalman filter demonstrates increased performance with binned spike rates, so a sliding window of 10 time samples was used to bin the neural data before input to the Kalman filter. Additionally, the LSTM decoder was trained on 30 online reaches after the offline training to increase robustness, a technique that has been shown to improve online decoding accuracy for machine learning models [44].

After offline training, the different decoding models performed 3 closed loop simulation experiments. The performance of each decoder is measured using average time to secure the target, a common metric for evaluating closed loop BMI performance [26, 27]. For each experiment, 10 simulations are performed with different neurons selected for disruption. The results in Fig. 7 are averaged across all 10 simulations. In the first experiment, 30 out of 46 input neurons were removed after 50 reaches. This simulates a loss of connection to neurons that is common in real BMI systems due to electrodes shifting or becoming damaged [25,50]. The continuous learning SNN was allowed to train using continuous learning for 30 subsequent reaches, while the other decoders were re-calibrated on neural data from the first 30 reaches after reassignment. Here, the SNN with continuous learning can learn without any pause in the usage of the BMI device whereas the BMI system has to be paused to allow calibration of the other decoders. The continuous learning SNN gets closest to a return to baseline performance, as seen in Fig. 7, with the largest performance drop seen by the BPTT SNN.

**Figure 7.**
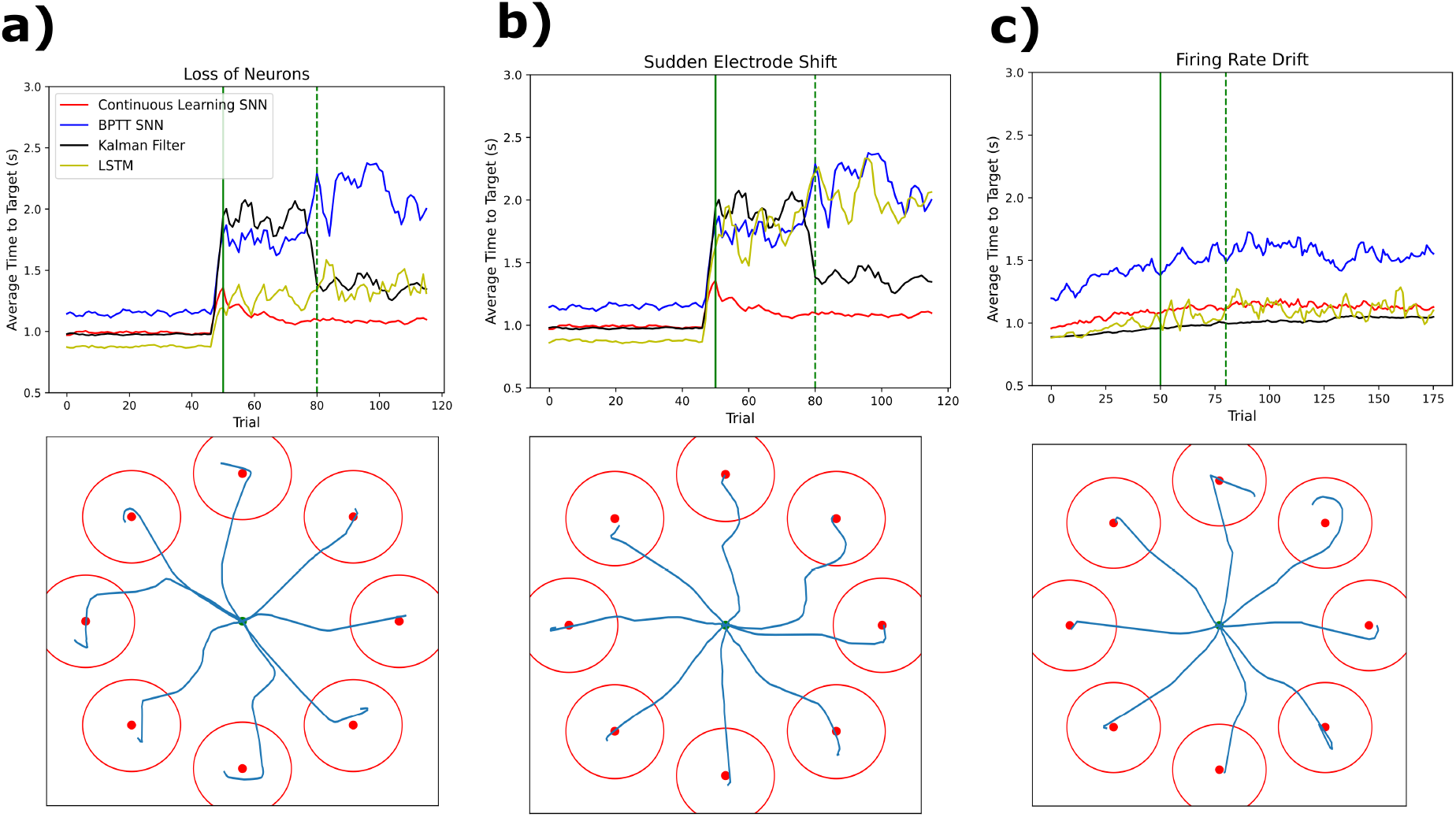
Plots of average time to target (top) and selected reaching paths (bottom) of simulated closed loop experiments testing three different kinds of neural disruption: neural dropout (a), changing kinematic mapping (b), and shifting firing rates (c). The top graphs show the average time required to secure the target for the continuous learning SNN (red), BPTT SNN (blue), ReFIT Kalman Filter (black) and LSTM decoder (yellow). The results are averaged across 10 different simulations, with a sliding window average of 4 reaches also applied to smooth out the results. The bottom graphs show the path taken by the continuous learning SNN decoder on eight example reach tasks after any neural disruption and retraining. The red circles around each target represent the window that the decoder has to stay inside for the reach to be considered successful. In (a) and (b), the solid green line represents the time at which the neural disruption occurred, while in (c), it represents the time at which firing rates stabilized. For all experiments, SNN continuous training starts at the solid green line and stops at the dotted green line. The other decoders are re-trained after the dotted green line using data from the 30 reaches performed between the two green lines.

In the second experiment, the preferred directions of 30 neurons were randomly reassigned. Although real neurons do not suddenly change behavior like this, it is common for electrodes used in implantable BMI to shift during use and start picking up data from a new set of neurons [25, 28], which can be simulated by changing the properties of the existing neurons. All decoders showed a large decrease in quality following neural reassignment. As with the previous experiment, the decoders are re-trained using 30 reaches after the simulated disruption. The continuous learning SNN was able to return to close to pre-disruption performance, while the performance of other decoders remains worse than pre-disruption levels. (Fig. 7). Additionally, the SNN training can be more easily implemented in a real BMI use setting; if a user notices a sudden decrease in quality, they can perform reaching tasks with continuous learning until quality improves. The other decoders cannot learn continuously, so to fix a performance drop a user must instead perform an arbitrary number of reaches and then wait for the decoder to be retrained on the new data.

In the last experiment, the maximum firing rates of the input neurons were allowed to change for the first 50 reaches. Real neurons have been shown to demonstrate changes in their firing rates over time [51], and degrading electrode quality can also cause apparent neural firing rates to decrease an an electrode ages. For this experiment, neurons were assigned a new “target” maximum firing rate between 0 and 30 spikes per second. The neurons approached the new firing rates exponentially with time constants chosen to be normally distributed with a mean of 30 seconds. After firing rate drift, the SNN trained continuously for 30 reaches, while the other decoders were re-trained after 30 reaches. All systems show a moderate decrease in performance which does not appear to improve after retraining, although the effect is most dramatic for the BPTT SNN. This suggests that although the continuous learning SNN decoder is fairly robust to changing firing rates, it is not capable of compensating for those changing rates by additional training.

#### 3.2.3. Closed Loop Learning

An additional simulation was performed to see if the SNN was capable of learning in a closed loop setting without prior offline training. An untrained SNN was trained online using continuous learning for 30 reaches, and then tested for an additional 70 reaches post training; Fig.8 shows the average time to secure the target. Here, online learning means that the output of the SNN was used to determine the cursor velocity from the beginning, even when the model parameters were completely randomized, and learning was performed by comparing the SNN output to the ideal direction of movement. As a result, the movement in the beginning will be uncontrolled and somewhat random; the goal is to show that the SNN can learn from this random movement and update its weights accordingly to allow more controlled movement. This is contrasted from the offline learning used to train the models in the subsection prior to disruption, where the cursor path is fixed and the output of the SNN is not actually used to control the cursor. In a real-world BMI experiment, the offline training would be performed by asking a subject to imagine moving a prosthetic, whereas online training would involve asking the person to actually move the prosthetic and performing weight updates to correct any errors in movement. The continuous learning SNN converged rapidly, achieving average times under 1.0 seconds (the minimum possible time to secure the target is 740 ms). For comparison, other untrained decoders were also subjected to a similar trial, including a ReFIT Kalman filter, LSTM, and another SNN. The first 30 reaches were conducted online using parameters that were randomized and constrained to a range of values that would produce reasonable outputs. After 30 reaches, the Kalman filter was fit to the neural data from the 30 training reaches, while the SNN and LSTM were trained on the 30 reaches using backpropagation. Unlike the continuous learning SNN, the other decoders were unable to learn from the online reaching tasks, and failed to reach the target within the 3 second limit for the majority of trials. This demonstrates that the continuous learning SNN is capable of learning entirely online, without a need for imagined offline reaches, which could make BMI calibration much easier for new users.

**Figure 8.**
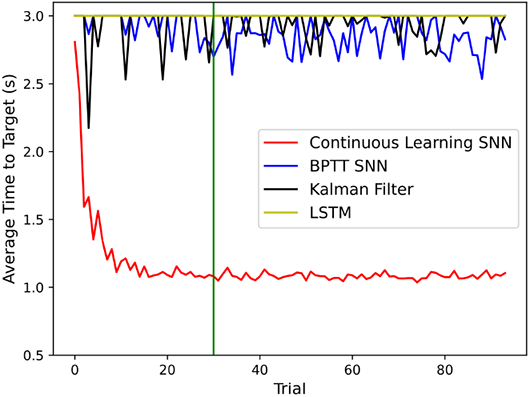
A plot showing the average time taken to reach the target for a continuous learning SNN, BPTT SNN, Kalman filter, and LSTM that received no prior training. The results shown are averaged across 10 different simulations. The output of each decoder is used to control the cursor during training, so the time to reach the target is not fixed, unlike offline learning. The continuous learning SNN learns rapidly, requiring only 30 reaches of online trials to achieve the same average time to target achieved by the decoders that were trained offline in the previous experiment. The other decoders are unable to learn from online trials, and typically fail to acquire the target.

## 4. Discussion and Conclusion

### 4.1. Significance

The continuous weight update mechanism for neural decoding SNNs presented in this paper has two significant applications for practical brain machine interfaces. First, it allows training on an arbitrary duration input with a fixed amount of memory. This makes it significantly easier to implement the algorithm in hardware, since there is no need for dynamic allocation of memory or introducing a hard limit on the input duration. One of the primary motivations for using SNNs is their power efficiency [23], which could enable safe implantation of a decoder directly inside a user’s brain, with no need for connection to an external device. In order to realize this power efficiency, a hardware implementation will be necessary, and this work takes an important step in that direction by introducing learning with a finite amount of memory. Future work should aim to implement the algorithm in a hardware system and measure its true power efficiency.

Additionally, the continuous nature of the weight update mechanisms presented here make the SNN ideal for use in a closed loop system. Adaptability is an important trait for online decoders, since implanted electrodes tend to shift over and degrade in quality over time, which can cause decreases in neural performance. The SNN presented here demonstrates an ability to maintain high performance after multiple different types of neural disruptions, including neural dropout, introduction of new neurons, and changing neural firing rates, with average time-to-target remaining within 20% of the pre-disruption baseline level after retraining for each disruption. The SNN can also train rapidly in an exclusively online setting, with no need for prior offline training, an ability not shared by other standard online decoders. This could reduce the difficulty of BMI calibration tasks for new users by removing the need for offline training in which the user must imagine a movement without physical feedback.

### 4.2. Limitations and Future Research

The proposed model does have some limitations that should be addressed in future work. It still requires more memory than the BPTT training algorithm in cases where the input duration is extremely short and the number of inputs is large, as shown in Subsection 3.1. Additionally, although the decoder enables faster adaptation to neural disruptions than existing methods, it still needs to be recalibrated with a sequence of designed reaching tasks to provide ground truth data, as do all other decoders examined in this paper. Future work should invest unsupervised calibration methods to allow algorithms to adapt with minimal disruption to the user.

Future research should also explore methods to optimize the SNN learning algorithm. Currently, the weight update algorithm is based on stochastic gradient descent. As shown in results, the continuous learning algorithm falls slightly short of the peak accuracy achieved by BPTT when trained offline with the Adam optimizer. Future work should investigate whether the accuracy of the continuous learning SNN can be improved by implementing more sophisticated optimization techniques, such as momentum.

Subsequent studies should also investigate the performance of the SNN in a real closed loop trial using animal model. Simulations have been demonstrated to be reliable predictors of actual closed loop performance, but a verification is still an important step before the algorithm is ready for use in an actual clinical care setting.

## Code

The code used in this paper, including the continuous learning SNN, BPTT SNN, and the code used to run the closed simulations, can be found at https://code.umd.edu/sshah389/spiking-based-brain-machine-interface.

